# Effect of ‘spent’ nucleotides on nonenzymatic RNA replication

**DOI:** 10.1101/2023.07.20.549979

**Authors:** Gauri M. Patki, Sudha Rajamani

## Abstract

Nonenzymatic template-directed replication would have been affected by co-solutes in a heterogenous prebiotic soup due to lack of enzymatic machinery. Unlike in contemporary biology, these reactions use chemically-activated nucleotides, which undergo rapid hydrolysis forming nucleoside monophosphates (‘spent’ monomers). These co-solutes cannot extend the primer but continue to base pair with the template, thereby interfering with replication. We therefore aimed to understand how a mixture of ‘spent’ ribonucleotides would affect nonenzymatic replication. We observed inhibition of replication in presence of the mixture, wherein predominant contribution came from the cognate Watson-Crick monomer, showing potential sequence dependence. Our study highlights how nonenzymatic RNA replication would have been directly affected by co-solutes, with ramifications for the emergence of functional polymers in an RNA World.

## Introduction

The RNA World hypothesis endeavours to provide an explication for the emergence, replication and evolution of genetic material in the absence of enzymes[1–3] during life’s origins. However, the efficiency of such nonenzymatic replication processes remains deficient. This contention about the efficiency is especially driven by two critical steps of replication (in addition to strand separation); selection of the correct nucleotide for incorporation against a templating base, and the proofreading mechanism that corrects for an incorrect incorporation. In the cellular milieu, these two steps are stringently facilitated by enzymes during enzymatic replication process. Thus, the error rates associated with enzymatic processes remain very low, ensuring faithful replication in extant biology[4]. Conversely, high error rates have been associated with nonenzymatic replication[5]. Although nonenzymatic replication raises concerns about the fidelity of replication, it has been shown that stalling after incorporation of a mismatched base pair could have intrinsically prevented ‘error catastrophe’[6]. Therefore, correct base pairing, which impinges on the appropriate stereochemistry of the extension terminus, is in itself a critical factor for determining fidelity of nonenzymatic replication[7].

Generally, nonenzymatic template-directed replication reactions have been studied by characterizing the primer extension process, using activated or nonactivated nucleoside monophosphates[8–12]. In a typical reaction, the template first forms a duplex with the primer, which is then followed by a nucleophilic attack of the 3’ end of the primer onto the incoming nucleotide, resulting in the extension of the primer by one nucleotide at a time[13]. Activated nucleotides bear a good leaving group, which makes the process of nonenzymatic primer extension feasible under ambient aqueous conditions, without invoking high temperatures or external catalysts[8]. However, these activated moieties are extremely unstable and prone to getting hydrolysed into the corresponding nucleoside 5’-monophosphates (NMPs). The NMPs thus formed, also called as ‘spent’ monomers, cannot extend the primer on their own under the aforementioned reaction conditions. However, the spent monomers possess an intact base pairing moiety, which allows them to continue to base pair with the templating base[14]. This affinity of the spent monomers for the templating base, can hinder the access of the activated nucleotides to the templating site, consequently altering the reaction kinetics of subsequent primer extensions[14, 15]. Thus, in addition to governing the fidelity of nonenzymatic replication, base pairing also factors importantly in potentially inhibiting the reaction in these aforesaid scenarios. Pertinently, NMPs would have been present as co-solutes in the prebiotic soup thereby underscoring the importance of systematically characterizing their inhibitory potential in prebiotic reactions. In the context of co-solutes, it has been previously reported that molecules such as lipids and polyethylene glycol (PEG) tend to reduce reaction rates[16]. Specifically, they have been shown to affect the rate of incorporation of imidazole-activated purine nucleotides against pyrimidine templating bases, in template-directed replication reactions. In these reactions, PEG was used as a nonspecific co-solute and a proxy for macromolecular crowders in the prebiotic soup as macromolecular crowding is known to affect several reaction rates[17–20].

Because prebiotic soup would have been a conglomerate of co-solutes, and as some co-solutes have been shown to affect the rate of enzyme free reactions, we set out to systematically assess the effect of a mixture of NMPs as co-solutes on nonenzymatic replication reactions. To the best of our knowledge, previous studies have looked at the inhibitory effect of only a single species of NMP on the primer extension process, at a given time. This was determined primarily by calculating the dissociation (and inhibitory) constants of the binding between the nonactivated nucleotide and the template in different sequence contexts[14, 15]. Moreover, a comprehensive study looking at the effect of a mixture of all four spent nucleotides on templated-replication in an all-RNA system involving ribonucleotides, RNA primer, albeit amino-terminal, and RNA templates is lacking. To address this lacuna, we investigated how the presence of a mixture of AMP, GMP, CMP and UMP (mixture of NMPs) would affect reaction rates, as they all would have been simultaneously present in a heterogenous prebiotic soup. This gains importance when accounting for the fact that a sophisticated replicating protein machinery was not present then. Therefore, the interference from these co-solutes due to their base pairing ability with the template would have directly influenced the reaction rates and their outcomes[15, 21]. Furthermore, a combination of intermolecular interactions, such as stacking, hydrogen bonding and secondary electrostatic interactions, is postulated to play a crucial role in determining the binding strength of a nucleotide to its templating base[15]. Such a complex interplay of interactions between the nucleotide and the template, makes it all the more important to characterize how the presence of a mixture of all the four spent nucleotides would have affected the rate of the reaction, given that each of them would have different binding strengths with the template, and thus different inhibitory potentials. All of this essentially raises two primary questions: (i) How would a mixture of NMPs affect nonenzymatic replication rates? (ii) How much would have the contribution of the individual components of the mixture been, towards this process?

Given the aforementioned reasons, we set out to understand the effect of a mixture of spent nucleotides on nonenzymatic templated-replication reactions in an all-RNA context. For this, we used a previously standardized reaction set up (Table 1,2; Methods and Materials)[16]. We observed that a 1:1:1:1 mixture of the four spent nucleotides (the NMP mixture), reduced the templated-replication rate significantly.

**Table 1:**
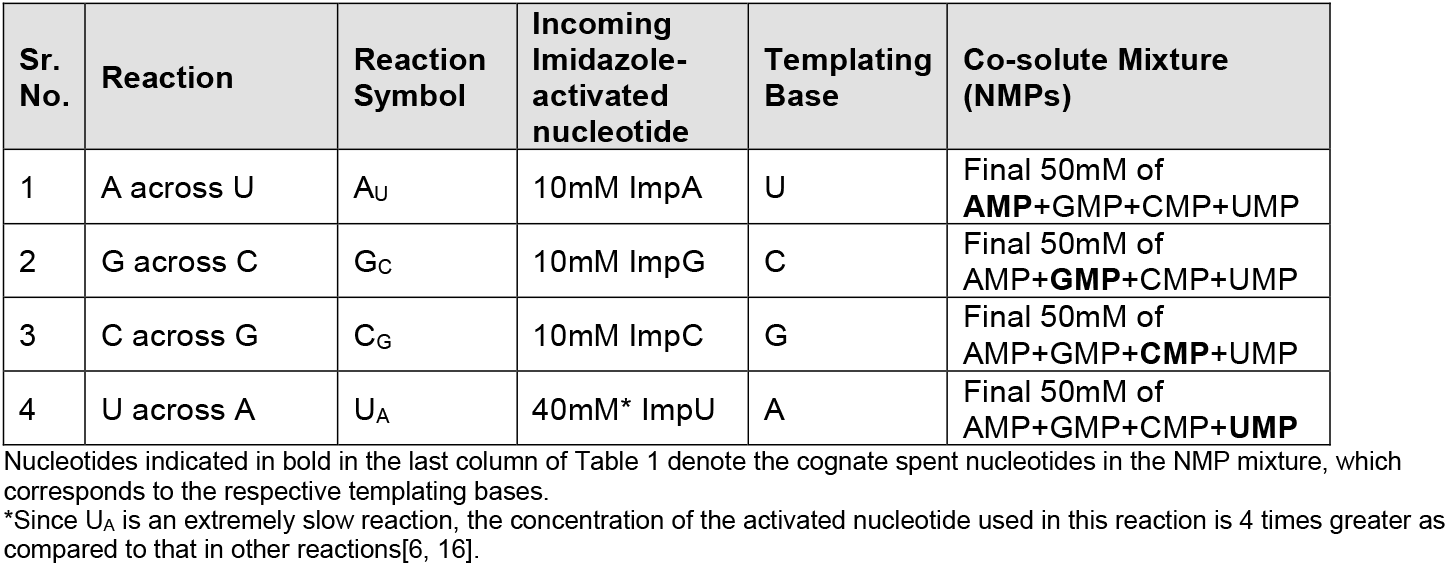
Reactions.

| Sr. No. | Reaction | Reaction Symbol | Incoming Imidazole-activated nucleotide | Templating Base | Co-solute Mixture (NMPs) |
| --- | --- | --- | --- | --- | --- |
| 1 | A across U | A<sub>U</sub> | 10mM ImpA | U | Final 50mM of **AMP**+GMP+CMP+UMP |
| 2 | G across C | G<sub>C</sub> | 10mM ImpG | C | Final 50mM of AMP+**GMP**+CMP+UMP |
| 3 | C across G | C<sub>G</sub> | 10mM ImpC | G | Final 50mM of AMP+GMP+**CMP**+UMP |
| 4 | U across A | U<sub>A</sub> | 40mM* ImpU | A | Final 50mM of AMP+GMP+CMP+**UMP** |

Nucleotides indicated in bold in the last column of Table 1 denote the cognate spent nucleotides in the NMP mixture, which corresponds to the respective templating bases.
\*Since U<sub>A</sub> is an extremely slow reaction, the concentration of the activated nucleotide used in this reaction is 4 times greater as compared to that in other reactions[6, 16].

**Table 2:**
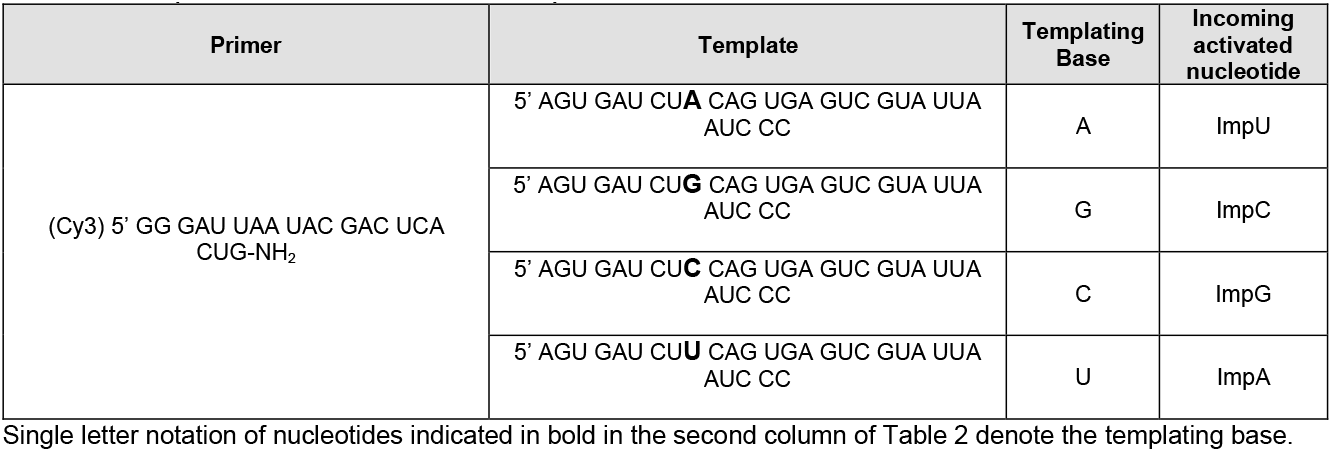
Sequences of Primer and Template.

Primarily, we observed that the spent nucleotides that prominently base pair with the templating base, were the ones that were mainly responsible for the effects that were observed. Furthermore, we observed predisposition of some reactions to inhibition, due to contribution from near-neighbour interactions and wobble base pairing that came from the spent nucleotides. Lastly, we conclude that the predominant factor responsible for reducing the reaction rate in the mixture, is the spent nucleotide that is the Watson-Crick base pairing partner of the templating base. In all, undertaking this study is pertinent to discerning how template-directed nonenzymatic replication of RNA in a putative RNA World, would have realistically panned out in the presence of a mixture of spent monomers in the vicinity. Moreover, this knowledge will have ramifications for characterizing prebiotically plausible mechanisms of ‘rescuing’ these replication reactions from ‘parasitic monomers’.

## Materials and Methods

### Materials

Nucleoside-5’-phosphorimidazolides (ImpNs) were purchased from GLSynthesis Inc. The RNA primer terminated with 3’-amino-2’,3’-dideoxynucleotide (Amino G primer) was acquired from Keck laboratory, Yale, USA and was gel purified before use. RNA templates were purchased from Sigma-Aldrich (Bangalore) and were used without any further purification. Sequences of the primer and template are mentioned in Table 1. Disodium salts of nucleoside-5’-monophosphates (5’-NMPdss) were purchased from Sigma-Aldrich (Bangalore). All other reagents were purchased from Sigma-Aldrich and were of alalytical grade.

### Methods

#### Reaction set-up and time points

1.3uM of respective template and 0.325 uM amino terminal primer (sequences as specified in the main text) were mixed and heated at 90°C for 5 minutes, after which the mixture was allowed to cool approximately for a minute. Tris (pH 7) and NaCl were added at final concentrations of 100mM and 200mM, respectively. NMPs were added in the form of a mixture of all the four NMPs or individually at a final concentration of either 50 mM or 12.5 mM, depending on the reaction. A stock of 5’-ImpN was made fresh and added to the reaction mixture at a final concentration of 10 mM. Final reaction volume was set to 10 ul. 1 ul of the reaction mixture was withdrawn at respective time points, put in TBE buffer containing 8 mM urea, and was frozen immediately at -35°C. Zero time point is withdrawn right after mixing the freshly made 5’ImpN with the reaction mixture.

#### Reference lane

As used in the previous study, respective control reactions are incubated separately for 1 hour and 1 ul of it is processed and loaded on the gel like the rest of the time points. These reactions are done separately and are only for the purpose of using for the reference positioning of the N and N+1 bands.

#### Quantification of the reaction

The amino primer bears Cyanine 3 (Cy3) fluorophore label at its 5’ end for its detection on the gel. Before loading the aforementioned time points on the gel, 15 times excess of an unlabelled competitive RNA, which is exactly identical in sequence with the amino primer, was added to the time points. The mixture was heated at 90 °C for 5 minutes, so as to separate template from the labelled primer. The individual time points were run on 20% urea PAGE. The gel was visualized by exploiting the Cy3 fluorescence at the 5’ terminal of the amino primer, which ensures detection of only primer bands. The extended primer (N+1) and the unextended primer (N) resolved into two separate bands on 20% denaturing PAGE, intensity of which was measured on ImageQuant L. Fraction of the extended product was calculated by following formula.

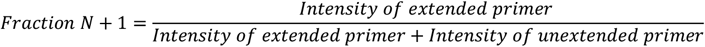

Reaction progress in the presence and absence of NMPs is compared by plotting Fraction N+1 against time. To determine the initial rate of each reaction, first few data points are fitted to a linear curve and slope of the same is determined.

#### Statistical Analysis

For all the statistical analysis, GraphPad Prism 9 was used. For a comparison between two reactions, Student’s unpaired two-tailed t-test was used. For a simultaneous comparison between individual reactions with control, Dunnett’s test was used. One Way ANOVA was used for multiple comparisons. For any given comparison between reaction populations, changes were considered statistically significant only for p-values <0.05.

## Results

### Effect of spent nucleotide mixture on the reaction rate of template-directed RNA replication

We carried out nonenzymatic replication reactions as mentioned in Table 1 with each reaction being assigned a reaction symbol as indicated in the table. The details of the primer template sequences and the activated imidazolides used in the reactions have been detailed in Table 2. In a typical reaction, a mixture of all the four spent nucleotides were added at a total concentration of 50mM to the reaction mixture (where the individual nucleotide’s concentration is 12.5mM). We hypothesized a reduction in the rates of all the four reactions in such a scenario because of two primary possibilities; (i) due to general crowding that is facilitated in these reactions by the presence of spent nucleotides as it could potentially affect the availability of the incoming activated nucleotide across the templating base (as was seen in an earlier study[16]) or, (ii) occurrence of specific (Watson-Crick) and/or nonspecific (non-Watson-Crick) interactions between the spent monomers and the templating base and other near-neighbour bases, all of which could retard the addition of the incoming nucleotide. For any given reaction (Table 1), the mixture of spent nucleotides would have one NMP that would be the Watson Crick base pairing partner of the templating base (indicated in bold), and it would henceforth be referred to as the ‘cognate spent monomer’ for the sake of brevity. Conversely, the rest of three spent monomers in the NMP mixture, are the non-Watson-Crick base pairing partners of the templating base, which would henceforth be referred to as the ‘non-cognate spent monomer’.

When the mixture of spent monomers was added as co-solutes in the replication reaction, a significant drop in the initial reaction rates was observed for A_U_, G_C_ and C_G_ reactions (Figure 1: (i)C, (ii)C and (iii)C). However, reaction U_A_ appeared to remain unaffected (Figure 1: (iv)C). It is pertinent to note that such an effect could be observed because of the higher ratio of ImpU:NMP mixture used in U_A_ reaction (4:5 of ImpU to NMP mixture) when compared to other reactions (where it is 1:5 of ImpN to NMP mixture). This result corroborated a previous observation wherein even 10 equivalents of dTMP showed only a modest reduction in the T across A reaction in the context of DNA[15]. In the same study, Richert and co-workers also quantified the binding strengths of the dNMPs to the corresponding Watson-Crick base pairing partners in the DNA template. This varied across the different templating bases, ranging from 10mM to 280mM. This prompted us to ask how much would the individual spent nucleotides in our RNA-based reactions (containing the NMP mixture) actually affect the reaction rates when added individually at a final concentration of 50mM in each of the above reactions. This would be important for understanding if the reduction is a result of a collective inhibitory effect of all the spent monomers in the reaction or is mainly coming only from specific monomers. Therefore, to delineate the contribution of the individual nucleotides to the reaction rate reduction (especially as was observed in the presence of the NMP mixture), we next carried out all the reactions in the presence of individual spent monomers.

**Figure 1:**
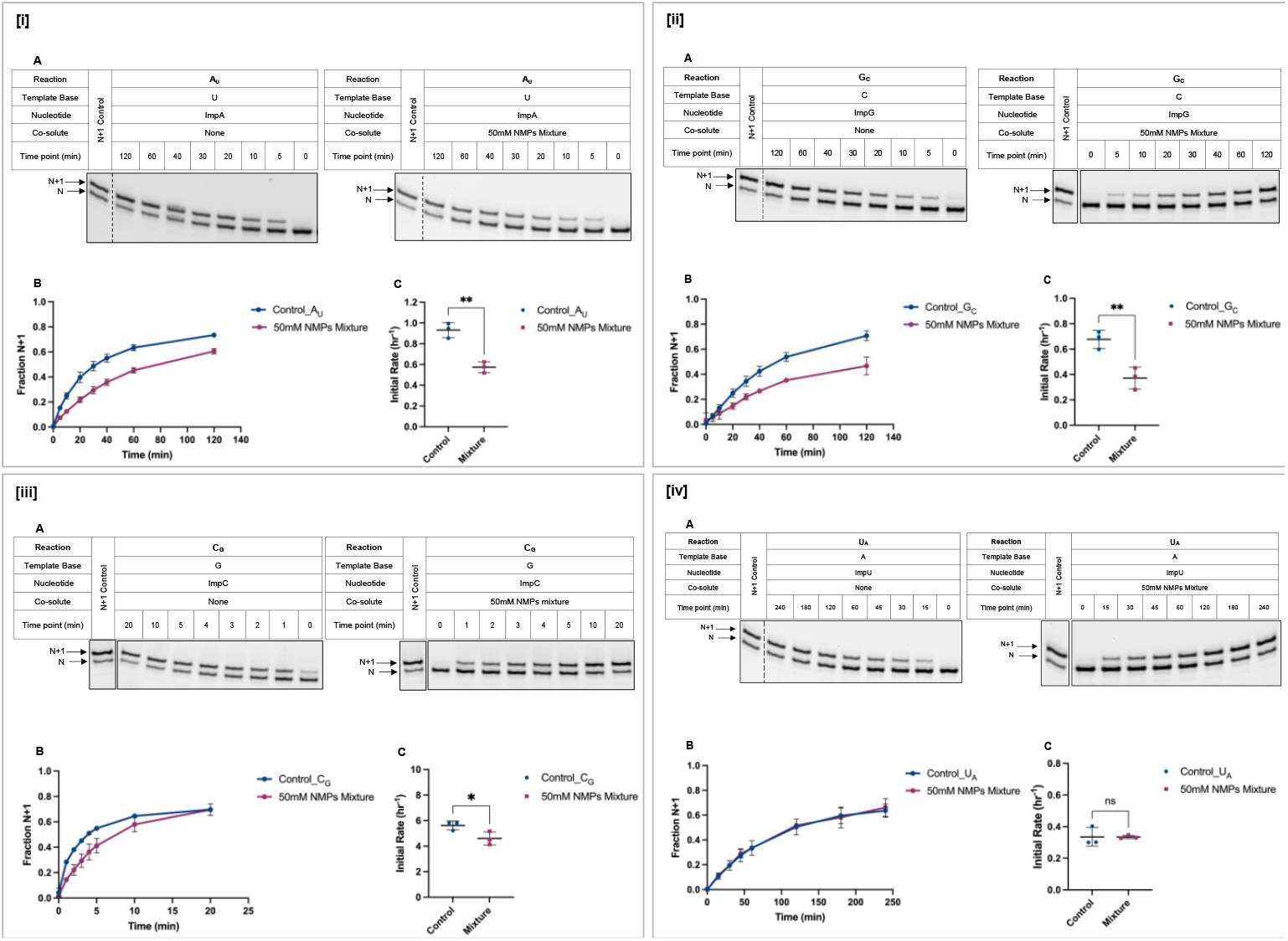
Effect of mixture of spent nucleotides on the rate of nonenzymatic template-directed reaction. Panels indicate **[i]** A_U_ reaction; **[ii]** G_C_ reaction; **[iii]** C_G_ reaction; **[iv]** U_A_ reaction, respectively. Subpanels indicate **A**. 20% denaturing gels of the relevant reaction in the absence (control) of co-solutes (left) and in the presence of 50mM NMPs mixture (right). Each gel image has a reference lane to represent the positions of the N and N+1 bands (Methods). Dotted line separates the reference lane from the adjacent lanes on the same gel. The reference lane is shown as a separate box when the sample was loaded a little away from the reaction of interest in the same gel (Full gel images have been included in SI; Figure S13-S18). **B**. Comparison of reaction progress between the control reaction and that with 50mM NMPs mixture for the reaction detailed in a given panel; **C**. Effect of presence of 50mM NMPs mixture (as co-solutes) on the initial reaction rate. For every reaction, the initial rate is calculated by fitting a linear curve to the first few time points before the reaction plateaus (methods and Figure S1-4); It is important to note that the values for initial rates are different for four different control reactions. Error bars represent standard deviation. N=3.

### Inhibition of nonenzymatic RNA replication by individual spent nucleotides

In the context of understanding the role of individual spent monomers as co-solutes in templated-replication reactions, it is reasonable to assume that the stronger the interaction between the spent monomer and the templating base, the more would be its ability to compete with the incoming activated nucleotide for the templating base. Consequently, the contribution of the cognate spent monomer in reducing the reaction rate would be greater when compared to others in the mixture. Indeed, it was clearly observed that the cognate spent monomer affects the rate of the reaction to a significantly greater extent than the non-cognate ones (Figure 2; A-C, Figure S5-S8). This set of data suggested that the rate reduction that we observed in the reactions detailed in Figure 1, could chiefly be because of the specific interactions of the cognate spent monomers with the templating base. This, in turn, could be causing hindrance in the interactions between the incoming activated nucleotide and the templating base.

**Figure 2:**
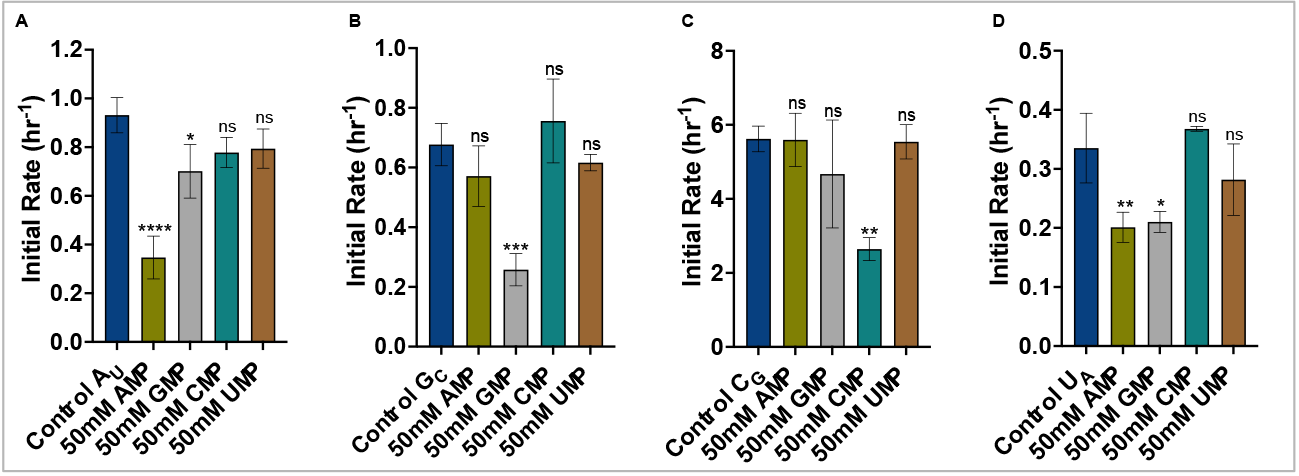
Effect of individual spent monomers on nonenzymatic template-directed replication reactions: **[A]** A_U_ reaction; **[B]** G_C_ reaction; **[C]** C_G_ reaction; **[D]** U_A_ reaction. The cognate spent monomers, AMP, GMP, and CMP in A, B and C, respectively, affect the progress of the reactions the most. These are indicated by green-, grey- and teal-coloured bars, respectively. In U_A_ reaction, the cognate spent monomer, UMP, did not show any effect on reaction rate (brown). Statistical analysis was done using Dunnett’s test, where means of all the reactions are compared to that of the control. Error bars represent standard deviation. N=3.

However, this scenario is not as straightforward as the aforementioned scenario indicates. Although the most significant decrease was seen in the case of the cognate spent monomer, two more cases showed remarkable effect too. Firstly, there was a significant decrease in the reaction rate in the presence of GMP as a co-solute in the A_U_ reaction (Figure 2A). It was evident from this rate reduction that GMP, a well-known wobble of U[21], does indeed compete with ImpA in the reaction, thereby significantly affecting the rate of this templated-replication reaction. Secondly, a statistically significant reduction in the U_A_ reaction rate was observed, when purines were present as co-solutes (Figure 2D). This effect could be attributed to the 3’ terminal guanine residue of the primer, as it can provide near-neighbour stacking interactions especially for the incoming purine nucleotides.

This effect stems from the fact that interactions other than direct base pairing are also crucial in determining reaction rates. These include near-neighbour interactions such as stacking of the incoming nucleotide with one of the bases in the primer, or with a downstream helper strand[22, 23]. In our study, we found that the slowest reaction U_A_, was affected significantly by spent purine nucleotides, which could be because of the potential of the terminal guanine residue of the primer to offer such stacking interactions to the incoming purine nucleotides. Other than the primer, the sequence of the template could also play a role in inducing effective stacking. This is especially relevant in our reaction as the templating base is followed by three pyrimidine residues (U, C, U), all of which can base pair with purine co-solutes and this allows for further stabilization via stacking interactions[24]. Since purine-purine stacking is stronger than purine-pyrimidine stacking, AMP and GMP potentially stacked better with the 3’-terminal guanine residue of the primer, resulting in the ability to successfully compete with the incoming ImpU, thereby reducing this reaction rate in our study. This further emphasizes the potential of near-neighbour interactions in determining nonenzymatic reaction rates, resulting in adverse effects especially in the presence of readily competing co-solutes. As for why the potential stacking effect was not seen to be prominent in the case of C_G_ reaction, this could result from the efficient base pairing capability of ImpC (three hydrogen bonds) with the template base G, which possibly could be compensating for the competition coming from the spent purine monomers.

Pertinently, Figure 2 shows that in isolation, the cognate spent monomer reduces the reaction rate even more when compared to the mixture (Figure 2 A-C vs Figure 1: (i)C-(iii)C). However, this observation could be attributed to the higher concentration of the cognate spent monomers that was used in the individual reactions, when compared to what was used in the mixtures. Since the mixture contains 1:1:1:1 ratio of all the 4 spent monomers, the concentration of the individual nucleotides in these scenarios is 12.5mM. Given this, we thought it was imperative to also understand whether the 12.5 mM concentration of the cognate spent monomer would also affect the rate by itself to the same extent, and how this compares to when it is present in the mixture (of NMPs). If this indeed turns out to be the case, it would be clear that the cognate spent monomer that was present in the mixture was predominantly responsible for inhibiting the reaction in the mixed spent nucleotides experiments.

### Cognate spent nucleotide chiefly affects the nonenzymatic replication reaction

When 12.5mM concentration of the cognate spent monomer was used in the reaction, we still observed a significant drop in the initial rate of the reactions when compared to the respective control reactions (Figure 3, S9-S11). We further compared the reaction rates involving 50mM and 12.5mM cognate spent monomers, versus the 50mM NMP mixture containing reactions (Figure 4: A-C). For this, we performed one-way ANOVA (multiple comparisons) to compare the means of each of the above (rates involving 50mM cognate spent monomer, 12.5mM cognate spent monomer, and 50mM NMP mixture) in A_U_, G_C_, and C_G_ reactions. Since there was no effect of the cognate spent monomer on the reaction rate in case of U_A_ (Figure 2D), we excluded the same from further analysis. In rest of the three cases, we observed a statistically non-significant difference in the reaction rates between 50mM NMPs mixture and 12.5mM cognate spent monomer (Figure 4: A-C). From this, it can be deciphered that the reduction that is being seen in the mixture, is predominantly being governed by the cognate spent monomer even in the mixed NMP containing reactions.

**Figure 3:**
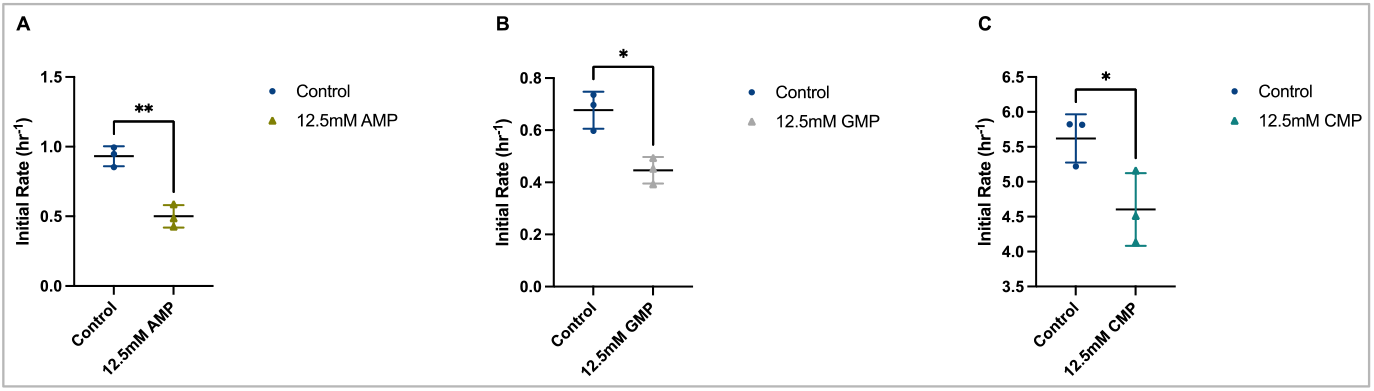
Effect of 12.5mM cognate spent monomer: **[A]** A_U_ reaction; **[B]** G_C_ reaction; **[C]** C_G_ reaction. Statistical analysis was done using Student’s unpaired two-tailed t-test. The linear fits for these are shown in Figure S9-S11. Error bars represent standard deviation. N=3

**Figure 4:**
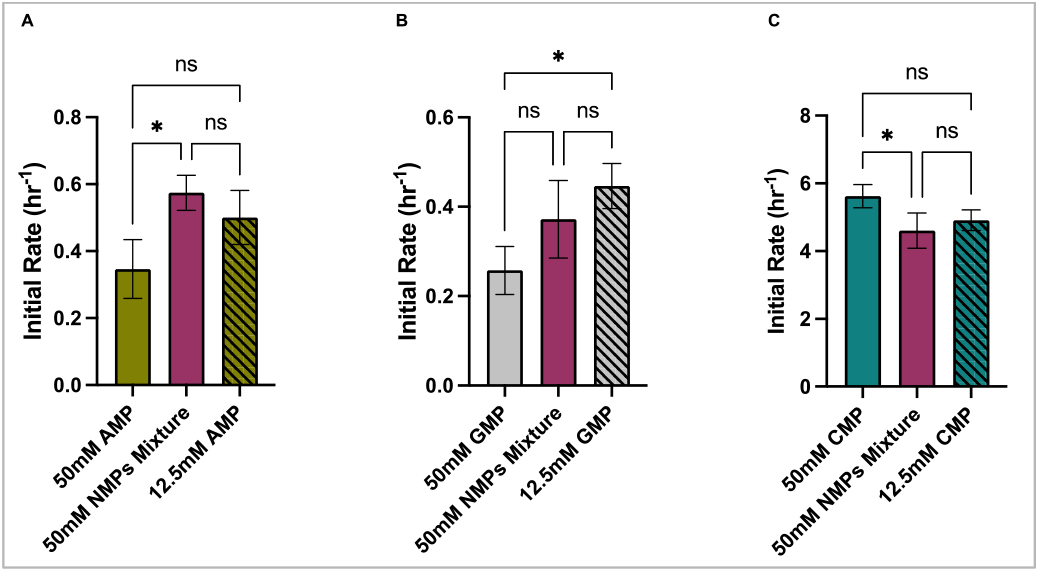
Identification of the determining factor for rate reduction: **[A]** A_U_ reaction; **[B]** G_C_ reaction; **[C]** C_G_ reaction. There is no statistically significant difference between initial rates of reactions containing 50mM NMP mixture and 12.5mM respective cognate spent monomer in all the three reactions, suggesting that the reduction that was observed in the case of mixed NMPs would have predominantly been brought about by the cognate spent nucleotide in the mixture. Statistical analysis was done using one-way ANOVA (multiple comparisons). Error bars represent standard deviation. N=3

## Discussion

Extant biology relies on transmission of genetic information, a process that is co-ordinated by a highly sophisticated enzyme machinery. This allows to distinguish between nucleotides and despite the presence of all four nucleotides (co-solutes) in the cellular milieu, faithful replication is still ensured with minimal errors. However, given the prebiotic heterogeneity and the absence of a complex enzymatic machinery, it becomes crucial to delineate how co-solutes in the prebiotic soup would have affected the rate of nonenzymatic templated-replication; one of the crucial milestones in a putative RNA World. In this context, we chose to look at spent nucleotides as co-solutes because of their availability in the prebiotic soup, their spontaneous formation as by-products from activated nucleotides, and their base pairing potential with the templating base. Also, we chose to study “all-RNA” reactions, which is the fundamental basis for the RNA World hypothesis. We addressed the effect of heterogeneity by checking the effect of a mixture of all four spent nucleotides on templated-replication.

Upon addition of 50mM NMP mixture to a given nonenzymatic template-directed replication reaction, the rate of the reaction significantly dropped. A_U_ showed 46.4% reduction in the initial rate while G_C_ showed 45.1% reduction (Figure S12) and C_G_ showed a comparatively lesser reduction at 20.6%. We observed that the rates of G_C_, C_G_ and U_A_ control reactions were slightly lesser when compared to an earlier study[16]. To address this, we systematically checked the hydrolysis profiles of all the ImpN monomers on HPLC (SI, Table S1). We observed hydrolysis in ImpG, ImpC, and ImpU monomers. We attribute the difference observed in the rates of control reactions when compared to the earlier study to this hydrolysis (as a result of which, the effective concentration of ImpN that went into the reaction was modestly lower than 10mM). Despite this observation, it is important to take into consideration that the concentration of the hydrolysed species was much lesser when compared to that of the co-solute that was externally added to the reactions, suggesting that the reduction in the rate would have been predominantly driven by the added co-solute. Although A_U_, G_C_, and C_G_ got affected, the effect seen in the case of U_A_, was negligible. It is also relevant to note that for an unambiguous comparison across various reaction sets, we wanted to keep the concentration of the co-solutes same across all the reactions (50mM), and thus, we did not change it for U_A_ despite the relatively higher concentration of ImpU that was used in the control reaction.

To understand if the effect observed is because of general crowding near the templating base, or if it is coming from other base specific interactions, we next checked how the individual spent nucleotide actually affected the reaction rate. We kept the ratio of activated:spent nucleotides at 1:5 ratio, excepting for U_A_ where the ratio was kept at 4:5. We observed that there was a direct correspondence between the propensity of base pairing of the spent nucleotide with the templating base, to the overall reduction observed in that reaction rate. Moreover, G-U wobble did seem to play a notable role in inhibiting the A_U_ reaction, once again signifying the importance of such non-Watson-Crick interactions even in the prebiotic context. In the U_A_ reaction, the presence of UMP did not show any effect on reaction rate. This could be because of the extremely poor potential of binding of UMP to the complementary templating base, such that the presence of it in excess was still inconsequential[21]. However, this same U_A_ reaction was observed to get affected by the presence of purines as co-solutes. And, this can be attributed to the potential stacking interactions between the spent purine monomers and the terminal guanine residue on the primer or the template, or both[24]. The exact role of template sequence in inhibiting the reaction rate, especially in the presence of spent purine nucleotides, can be confirmed by undertaking a detailed study to characterize the effect of mutations downstream of the templating base, or by having short sequence elements occupying those positions[22].

In all, observations from our study suggest the following about spent nucleotides as co-solutes: (i) The co-solute’s concentration and affinity towards the templating base determines the extent to which the rate would be affected (ii) Contribution from G-U wobble pairing in inhibiting the reaction is substantial (iii) Sequence of the primer and template, and consequently the near-neighbour interactions that can be facilitated, can determine how fast the reaction would proceed in the presence of co-solutes. This is significant as it indicates that the dependence of a spent monomer’s inhibitory action on a sequence of RNA, would have affected the replication of some sequences more readily than that of others. This would have had a direct implication for the evolution of the prebiotic RNA sequence space in the absence of enzymatic machinery. Specifically, in the absence of an enzymatic translation machinery, ‘single nucleotide translation’ is proposed to have happened, for which base-pairing of the aminoacylated nucleotide with the template is an essential requirement[25]. Therefore, spent nucleotides available in the vicinity could have had an inhibitory or regulatory role on these reactions as they would have directly competed for base-pairing with the templating base in these scenarios. Furthermore, replication happening inside phase separated systems containing nucleoside triphosphates (NTPs) as anionic moieties[26, 27] could also get affected, as the NTPs in these systems could potentially interact with the template and hinder the access of the activated nucleotides to the templating base.

Nevertheless, having a prebiotically plausible ‘scrutinizer’ may have saved such crucial reactions from ‘parasitic’ co-solutes. In this context, overcoming inhibition by removing the spent nucleotides from the reaction mixture has been studied[14, 28]. Here, we propose that replication of RNA inside fatty acid-based vesicles could have acted as a putative mechanism to overcome inhibition of spent nucleotides.[29, 30] The inherent characteristic of prebiotically relevant protocellular membranes could have, in principle, allowed for the selective diffusion of imidazole activated nucleotides over the spent monomers[29]. This is because spent nucleotides are charged molecules while imidazole activated nucleotides are not charged at neutral pH. This would have directly impinged on the permeability of the activated nucleotides across the bilayer, as the charge difference could have resulted in pertinent monomers being selectively favoured for replication over inhibition. This possibility is currently being investigated systematically by us. Significantly, this would have also depended on the nature of the protocellular membrane, thereby providing another selection pressure that could have also helped in potentially shaping the prebiotic amphiphilic landscape on the early Earth. Discerning the aforementioned aspects could shed light on whether the lipid boundary (and the type thereof) could have also saved replication reactions from spent monomers present in the vicinity, in addition to protecting them from parasitic sequences and metabolites. Taken together, our study comprehensively exemplifies how chemical diversity of the prebiotic soup, especially in the context of the concentration and affinity of its constituents towards templating bases, could have directly affected the error rates of nonenzymatic template-directed replication reactions.

## Acknowledgements

This research was supported by grants from the Science and Engineering Research Board (SERB), Department of Science and Technology, Government of India [CRG/2021/001851] and IISER Pune. The authors wish to extend their special thanks to Dr. Ramana Athreya for his help with statistical analysis, and to Dr. Shikha Dagar for her helpful discussions regarding the work. Thanks also to Kushan Lahiri for contributing towards Fig 1(iii) and Fig 2(C). G.M.P. also thanks the Department of Biotechnology, Government of India for fellowship support.

## Author Contributions

G.M.P. and S.R. designed the experiments while G.M.P. performed the experiments. G.M.P. and S.R. analysed the data. G.M.P. and S.R. wrote the manuscript.

## References

1. Gilbert, W. (1986). Origin of life: The RNA world. Nature, 319(6055), 618–618. https://doi.org/10.1038/319618a0

2. Robertson, M. P., & Joyce, G. F. (2012). The origins of the RNA world. Cold Spring Harbor Perspectives in Biology, 4(5), a003608. https://doi.org/10.1101/cshperspect.a003608

3. Szostak, J. W. (2012). The eightfold path to non-enzymatic RNA replication. Journal of Systems Chemistry, 3(1), 2. https://doi.org/10.1186/1759-2208-3-2

4. Loeb, L. A., & Kunkel, T. A. (n.d.). Fidelity of DNA Synthesis, 31.

5. Leu, K., Kervio, E., Obermayer, B., Turk-MacLeod, R. M., Yuan, C., Luevano, J.-M., … Chen, I. A. (2013). Cascade of Reduced Speed and Accuracy after Errors in Enzyme-Free Copying of Nucleic Acid Sequences. Journal of the American Chemical Society, 135(1), 354–366. https://doi.org/10.1021/ja3095558

6. Rajamani, S., Ichida, J. K., Antal, T., Treco, D. A., Leu, K., Nowak, M. A., … Chen, I. A. (2010). Effect of Stalling after Mismatches on the Error Catastrophe in Nonenzymatic Nucleic Acid Replication. Journal of the American Chemical Society, 132(16), 5880–5885. https://doi.org/10.1021/ja100780p

7. Blain, J. C., & Szostak, J. W. (2014). Progress Toward Synthetic Cells. Annual Review of Biochemistry, 83(1), 615–640. https://doi.org/10.1146/annurev-biochem-080411-124036

8. Leslie E. O. (2004). Prebiotic Chemistry and the Origin of the RNA World. Critical Reviews in Biochemistry and Molecular Biology, 39(2), 99–123. https://doi.org/10.1080/10409230490460765

9. Kervio, E., Sosson, M., & Richert, C. (2016). The effect of leaving groups on binding and reactivity in enzyme-free copying of DNA and RNA. Nucleic Acids Research, 44(12), 5504–5514. https://doi.org/10.1093/nar/gkw476

10. Prabahar, K. J., & Ferris, J. P. (1996). Adenine derivatives as phosphate activating groups for prebiotic RNA synthesis. Origins of Life and Evolution of the Biosphere, 26(3–5), 251–252. https://doi.org/10.1007/BF02459740

11. Bapat, N. V., & Rajamani, S. (2018). Templated replication (or lack thereof) under prebiotically pertinent conditions. Scientific Reports, 8(1), 15032. https://doi.org/10.1038/s41598-018-33157-9

12. Dagar, S., Sarkar, S., & Rajamani, S. (2023). Nonenzymatic Template-Directed Primer Extension Using 2′-3′ Cyclic Nucleotides Under Wet-Dry Cycles. Origins of Life and Evolution of Biospheres. https://doi.org/10.1007/s11084-023-09636-z

13. Walton, T., Zhang, W., Li, L., Tam, C. P., & Szostak, J. W. (2019). The Mechanism of Nonenzymatic Template Copying with Imidazole-Activated Nucleotides. Angewandte Chemie International Edition, 58(32), 10812–10819. https://doi.org/10.1002/anie.201902050

14. Deck, C., Jauker, M., & Richert, C. (2011). Efficient enzyme-free copying of all four nucleobases templated by immobilized RNA. Nature Chemistry, 3(8), 603–608. https://doi.org/10.1038/nchem.1086

15. Kervio, E., Claasen, B., Steiner, U. E., & Richert, C. (2014). The strength of the template effect attracting nucleotides to naked DNA. Nucleic Acids Research, 42(11), 7409–7420. https://doi.org/10.1093/nar/gku314

16. Bapat, N. V., & Rajamani, S. (2015). Effect of Co-solutes on Template-Directed Nonenzymatic Replication of Nucleic Acids. Journal of Molecular Evolution, 81(3), 72–80. https://doi.org/10.1007/s00239-015-9700-1

17. Kilburn, D., Roh, J. H., Behrouzi, R., Briber, R. M., & Woodson, S. A. (2013). Crowders Perturb the Entropy of RNA Energy Landscapes to Favor Folding. Journal of the American Chemical Society, 135(27), 10055–10063. https://doi.org/10.1021/ja4030098

18. Nakano, S., Miyoshi, D., & Sugimoto, N. (2014). Effects of Molecular Crowding on the Structures, Interactions, and Functions of Nucleic Acids. Chemical Reviews, 114(5), 2733–2758. https://doi.org/10.1021/cr400113m

19. Desai, R., Kilburn, D., Lee, H.-T., & Woodson, S. A. (2014). Increased Ribozyme Activity in Crowded Solutions*. Journal of Biological Chemistry, 289(5), 2972–2977. https://doi.org/10.1074/jbc.M113.527861

20. Ellis, R. J. (2001). Macromolecular crowding: obvious but underappreciated. Trends in Biochemical Sciences, 26(10), 597–604. https://doi.org/10.1016/S0968-0004(01)01938-7

21. Izgu, E. C., Fahrenbach, A. C., Zhang, N., Li, L., Zhang, W., Larsen, A. T., … Szostak, J. W. (2015). Uncovering the Thermodynamics of Monomer Binding for RNA Replication. Journal of the American Chemical Society, 137(19), 6373–6382. https://doi.org/10.1021/jacs.5b02707

22. Vogel, S. R., Deck, C., & Richert, C. (2005). Accelerating chemical replication steps of RNA involving activated ribonucleotides and downstream-binding elements. Chemical Communications, (39), 4922–4924. https://doi.org/10.1039/B510775J

23. Hagenbuch, P., Kervio, E., Hochgesand, A., Plutowski, U., & Richert, C. (2005). Chemical Primer Extension: Efficiently Determining Single Nucleotides in DNA. Angewandte Chemie, 117(40), 6746–6750. https://doi.org/10.1002/ange.200501794

24. Kanavarioti, A., Bernasconi, C. F., & Baird, E. E. (1998). Effects of Monomer and Template Concentration on the Kinetics of Nonenzymatic Template-Directed Oligoguanylate Synthesis. Journal of the American Chemical Society, 120(34), 8575–8581. https://doi.org/10.1021/ja9807237

25. Jash, B., Tremmel, P., Jovanovic, D., & Richert, C. (2021). Single nucleotide translation without ribosomes. Nature Chemistry, 13(8), 751–757. https://doi.org/10.1038/s41557-021-00749-4

26. Nakashima, K. K., van Haren, M. H. I., André, A. A. M., Robu, I., & Spruijt, E. (2021). Active coacervate droplets are protocells that grow and resist Ostwald ripening. Nature Communications, 12(1), 3819. https://doi.org/10.1038/s41467-021-24111-x

27. Liu, Z., Chen, J., Bai, Q., Lin, Y., & Liang, D. (2022). Coacervate Formed by an ATP-Binding Aptamer and Its Dynamic Behavior under Nonequilibrium Conditions. Langmuir, 38(20), 6425–6434. https://doi.org/10.1021/acs.langmuir.2c00580

28. Adamala, K., & Szostak, J. W. (2013). Non-enzymatic template-directed RNA synthesis inside model protocells. Science (New York, N.Y.), 342(6162), 1098–1100. https://doi.org/10.1126/science.1241888

29. Mansy, S. S., Schrum, J. P., Krishnamurthy, M., Tobé, S., Treco, D. A., & Szostak, J. W. (2008). Template-directed synthesis of a genetic polymer in a model protocell. Nature, 454(7200), 122–125. https://doi.org/10.1038/nature07018

30. Jin, L., Kamat, N. P., Jena, S., & Szostak, J. W. (2018). Fatty Acid/Phospholipid Blended Membranes: A Potential Intermediate State in Protocellular Evolution. Small, 14(15), 1704077. https://doi.org/10.1002/smll.201704077

