## Supplementary File for "Effect of ‘spent’ nucleotides on nonenzymatic RNA replication"

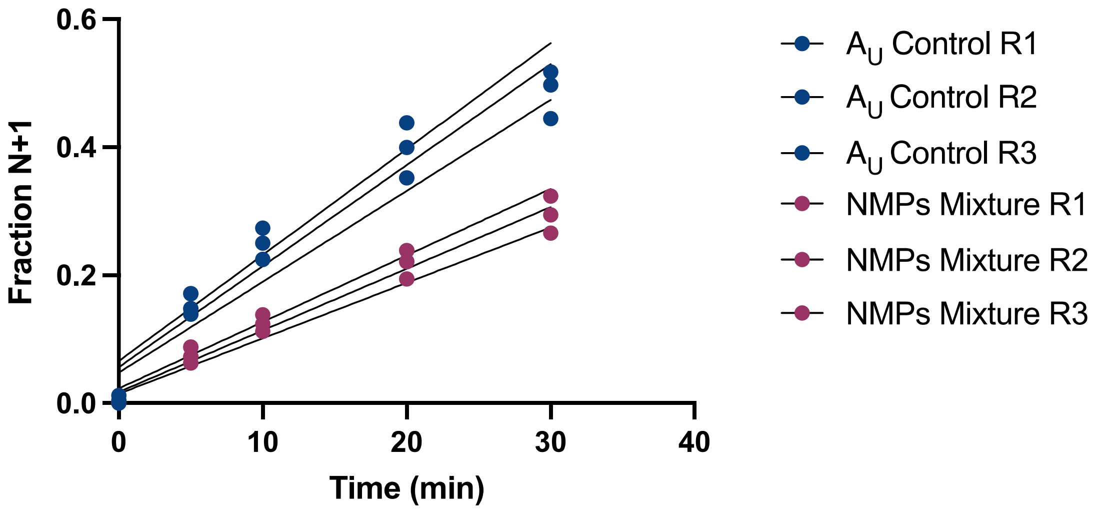


**Figure S1**: Linear fits for three replicates of initial rate calculations of $A_U$ control reaction and that of the 50mM NMPs Mixture.

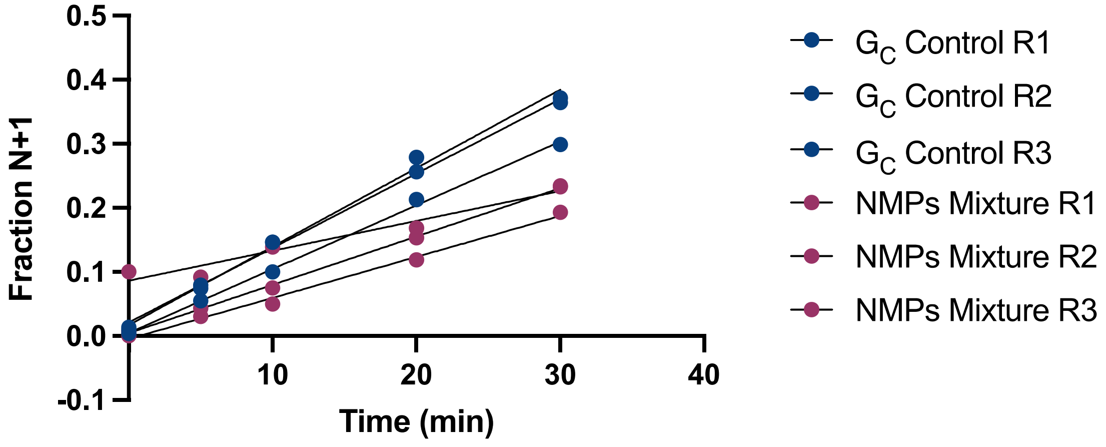


**Figure S2**: Linear fits for three replicates of initial rate calculations of $G_C$ control reaction and that of the 50mM NMPs Mixture.

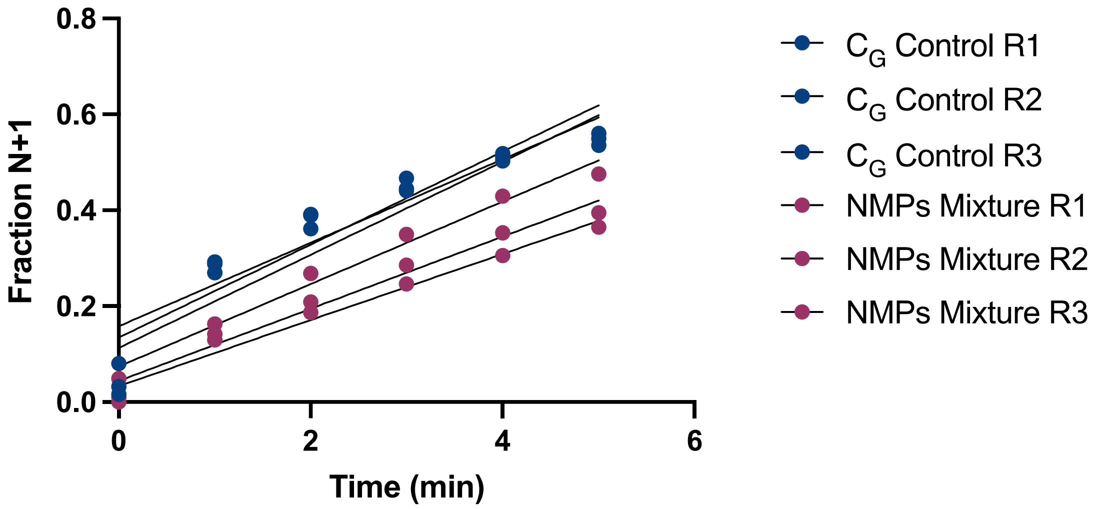


**Figure S3**: Linear fits for three replicates of initial rate calculations of $C_G$ control reaction and that of the 50mM NMPs Mixture.

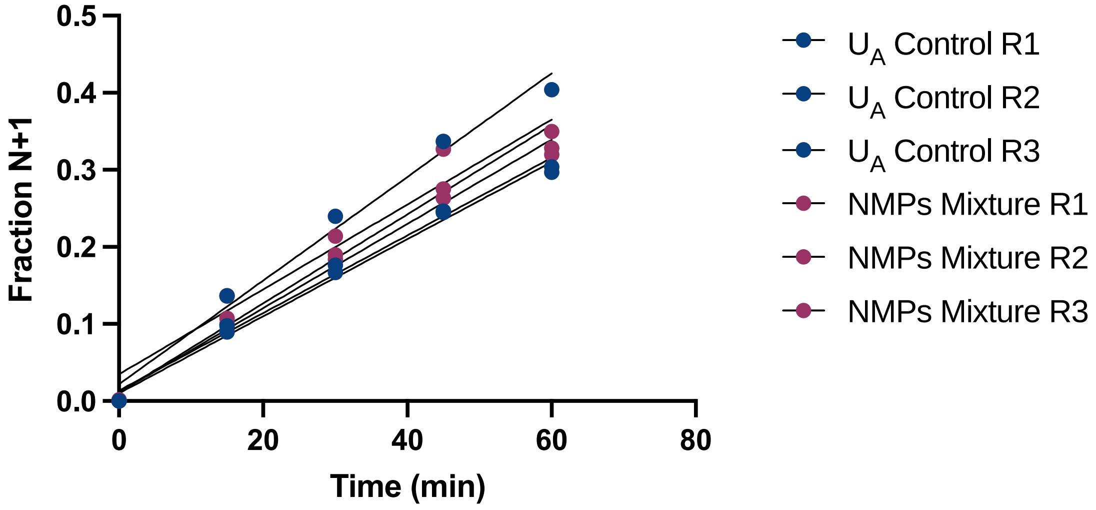


**Figure S4**: Linear fits for three replicates of initial rate calculations of $U_A$ control reaction and that of the 50mM NMPs Mixture.

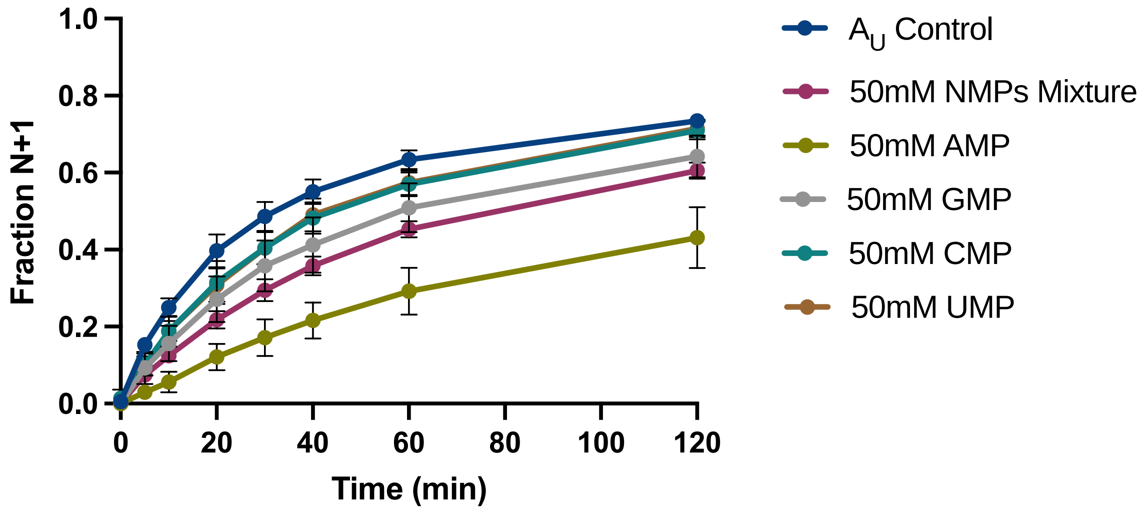


**Figure S5**: Reaction progress of $A_U$ reaction in the presence of various co-solutes.

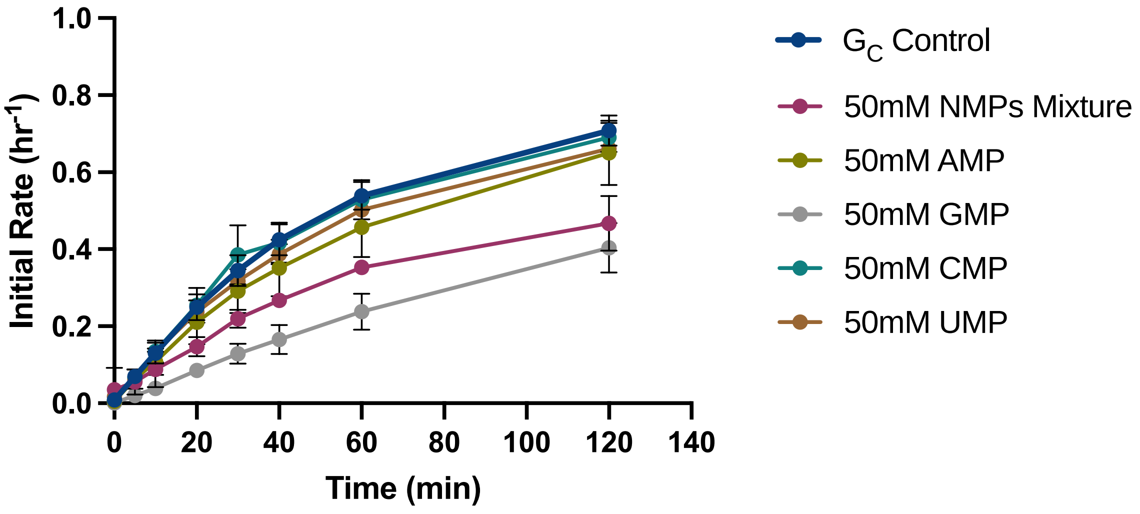


**Figure S6**: Reaction progress of $G_C$ reaction in the presence of various co-solutes.

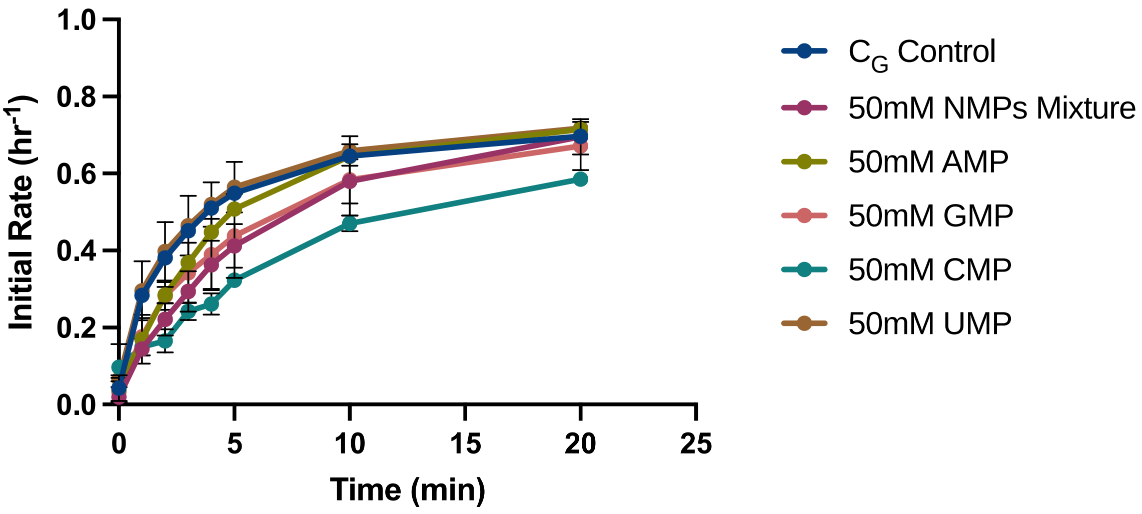


**Figure S7**: Reaction progress of $C_G$ reaction in the presence of various co-solutes.

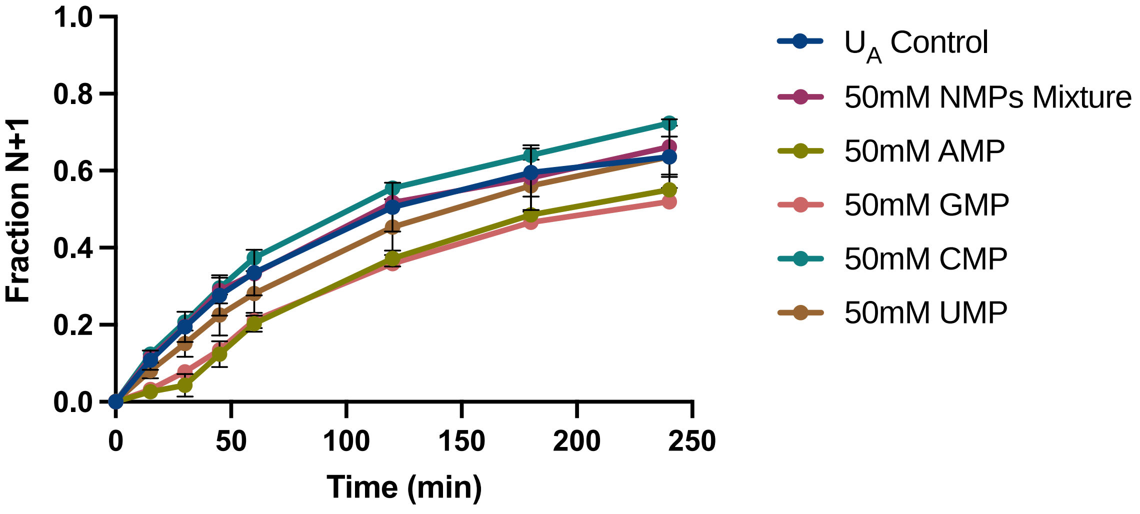


**Figure S8**: Reaction progress of $U_A$ reaction in the presence of various co-solutes.

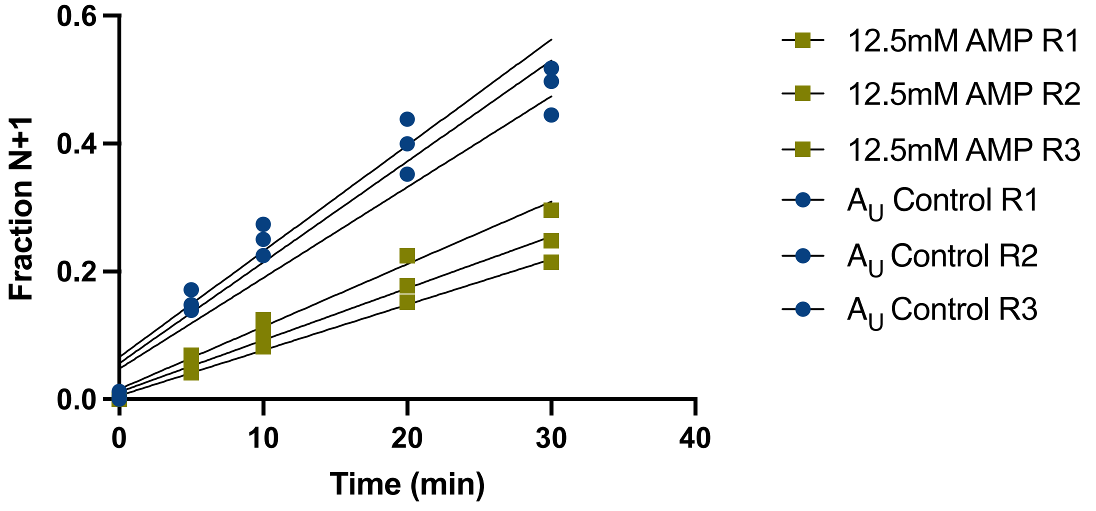


**Figure S9**: Linear fits for three replicates for initial rate calculations of $A_U$ control reaction and that of the 12.5mM NMPs Mixture.

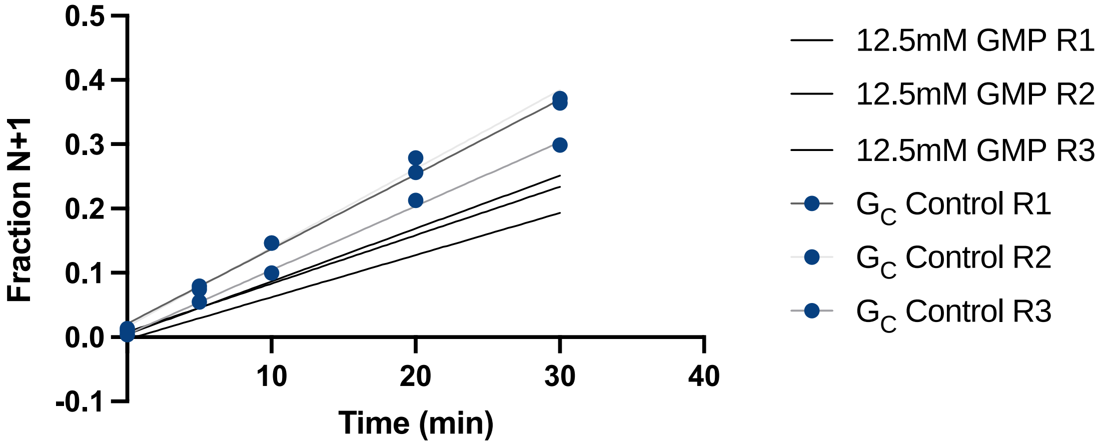


**Figure S10**: Linear fits for three replicates of initial rate calculations of $G_C$ control reaction and that of the12.5mM NMPs Mixture.

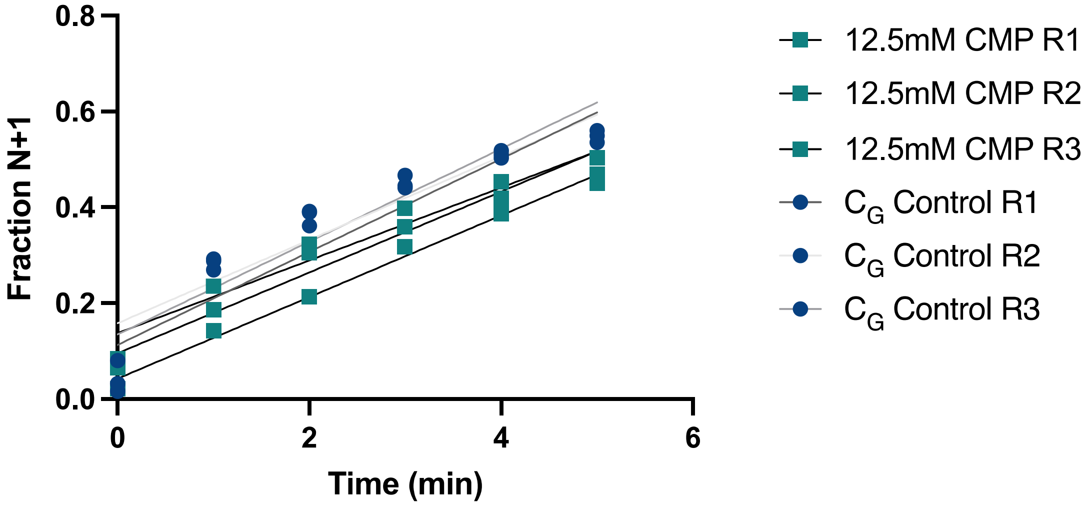


**Figure S11**: Linear fits for three replicates of initial rate calculations of $C_G$ control reaction and that of the 12.5mM NMPs Mixture.

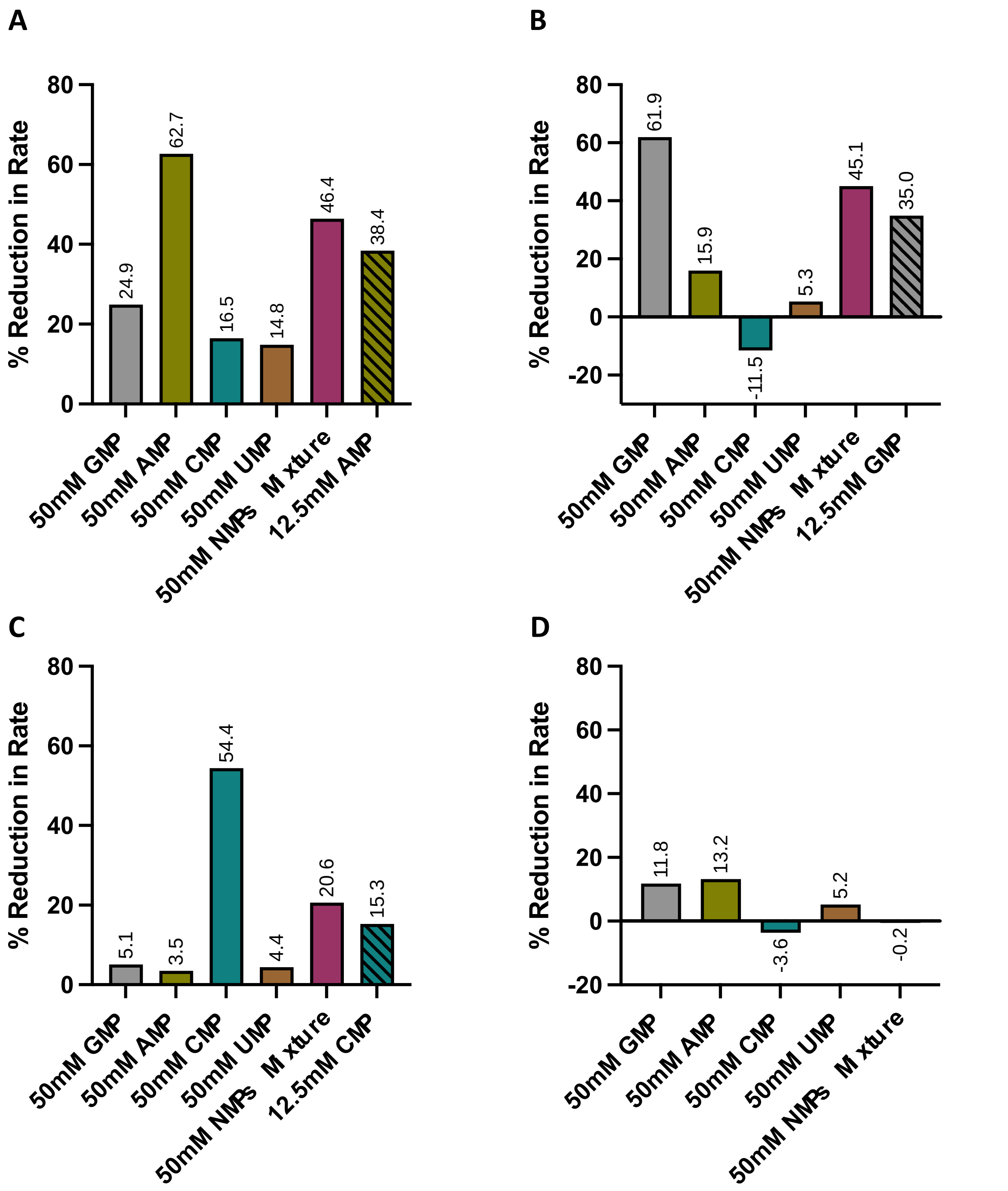


**Figure S12**: Percentage reduction in the initial rate of $A_U$ (**A**), $G_C$ (**B**), $C_G$(**C**), $U_A$ (**D**) reactions in the presence of various co-solutes (based on rates obtained for three reaction replicates, as indicated in above figures). The negative values that are observed in $G_C$ and $U_A$ reactions fall within the error range of the respective controls (Figure 2B, 2D and 1(iv)C), and do not affect the interpretation of the results.

#### Uncropped Original Full Gel Images used in the main text

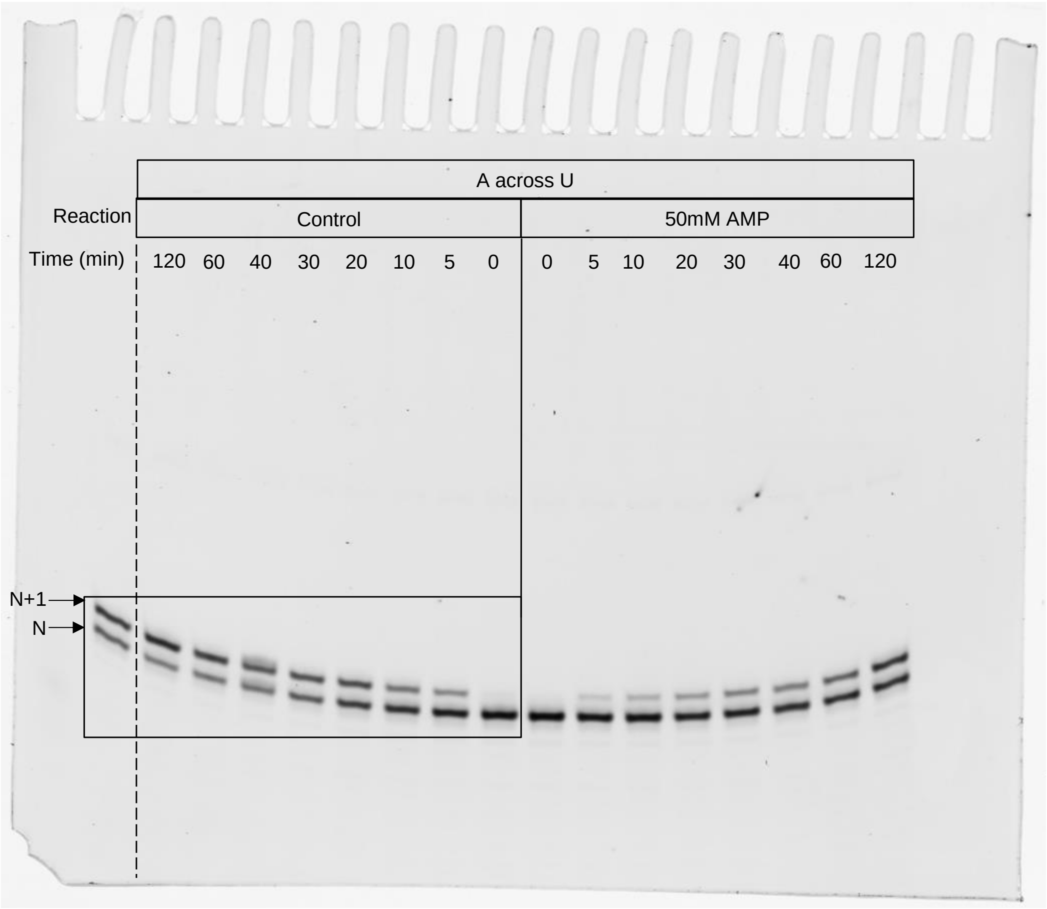


**Figure S13**: Box represents the cropped part of $A_U$ reaction shown in Figure 1(i)A in the main text.

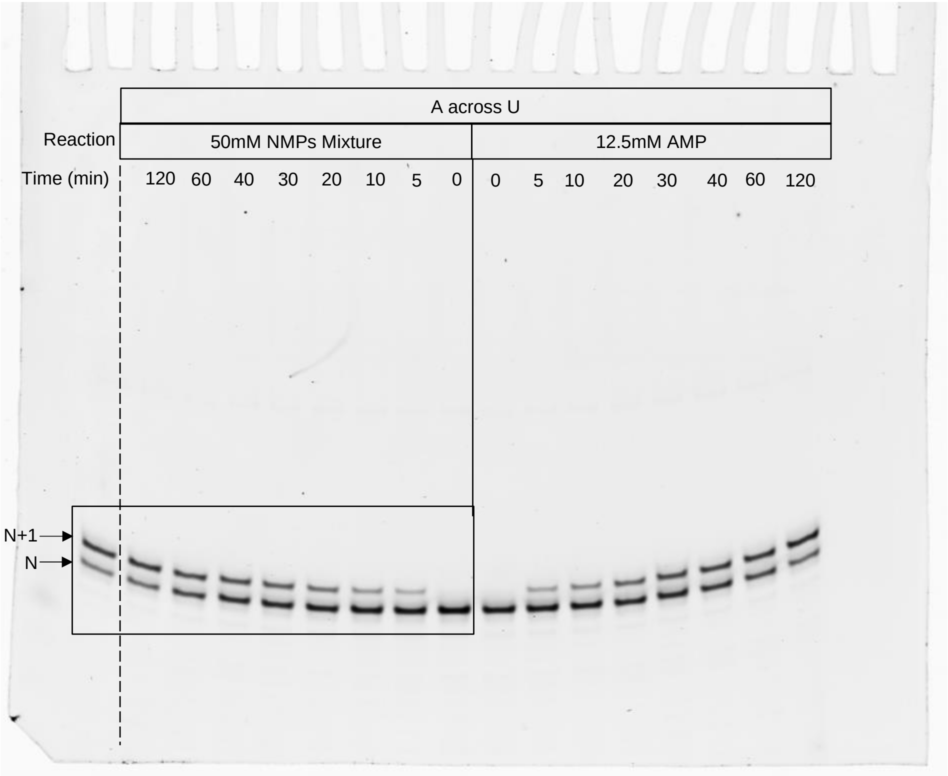


**Figure S14**: Box represents the cropped part of $A_U$ reaction shown in Figure 1(i)A in the main text.

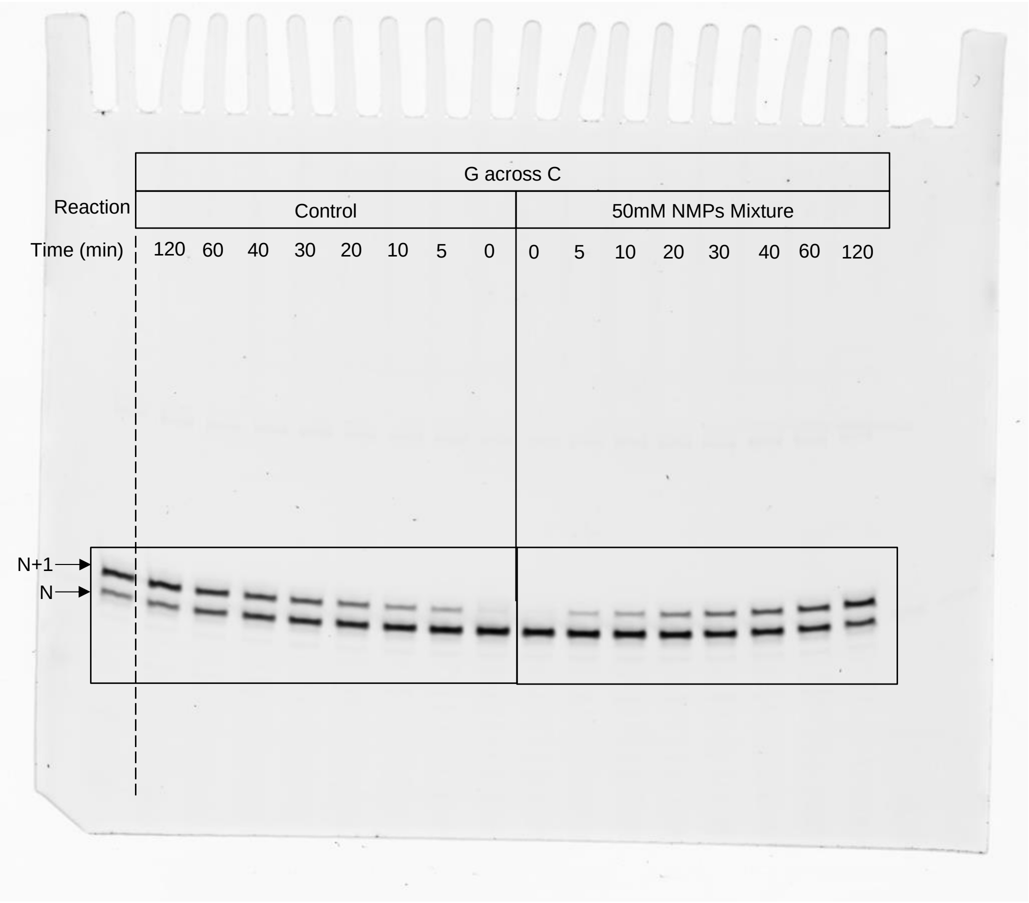


Figure S15: Boxes represent the cropped part of $G_C$ reaction shown in Figure 1(ii)A in the main text

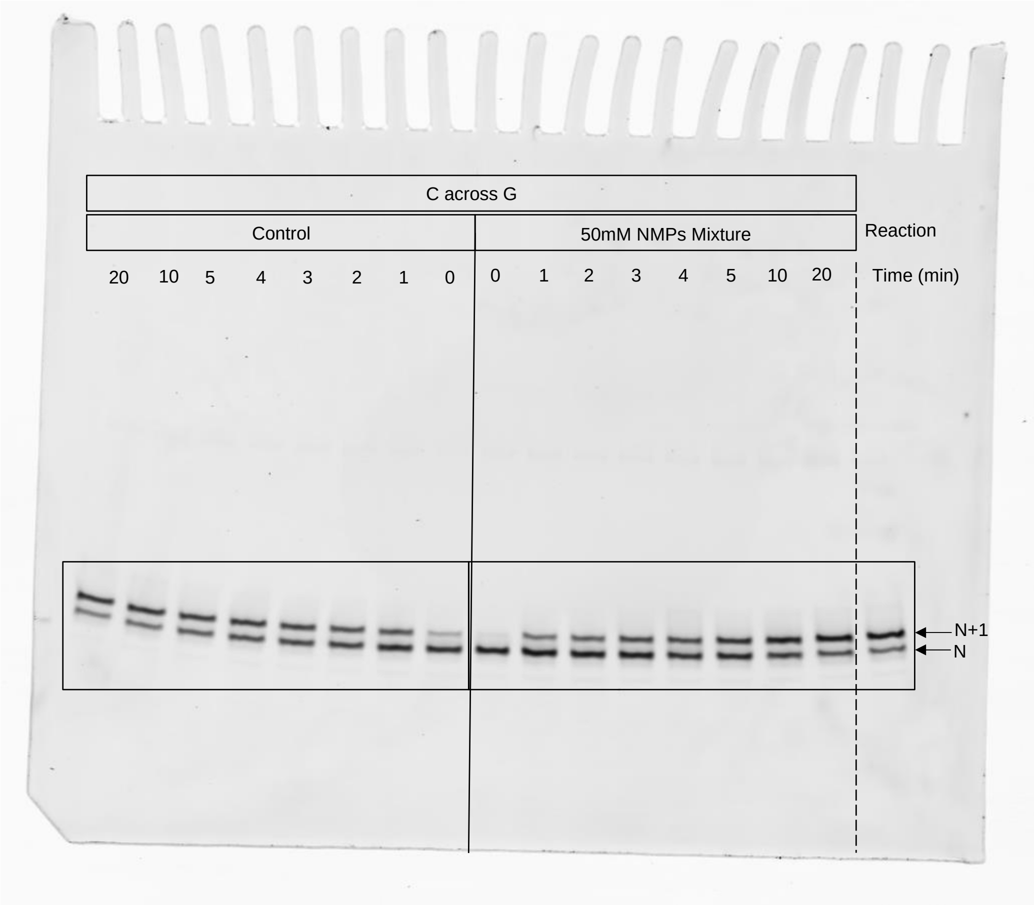


Figure S16: Boxes represent the cropped part of $C_G$ reaction shown in Figure 1(iii)A in the main text.

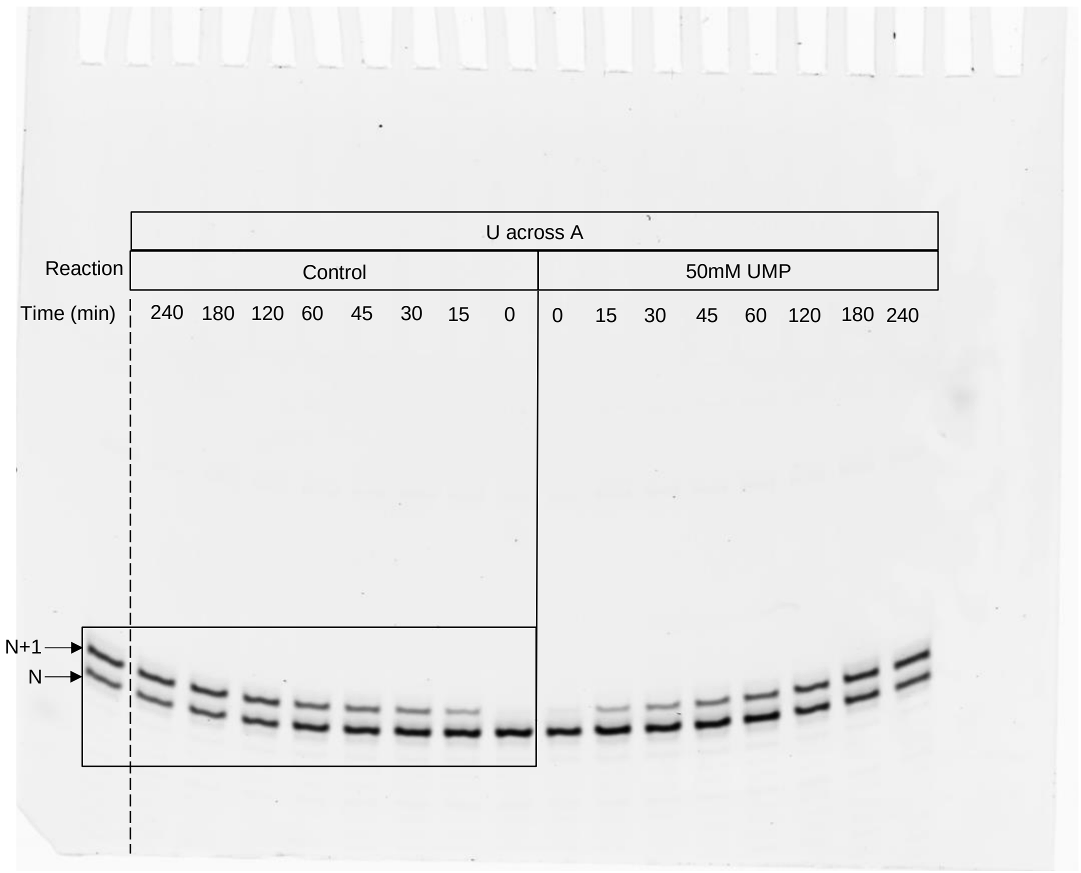


**Figure S17**: Box represents the cropped part of $U_A$ reaction shown in Figure 1(iv)A in the main text.

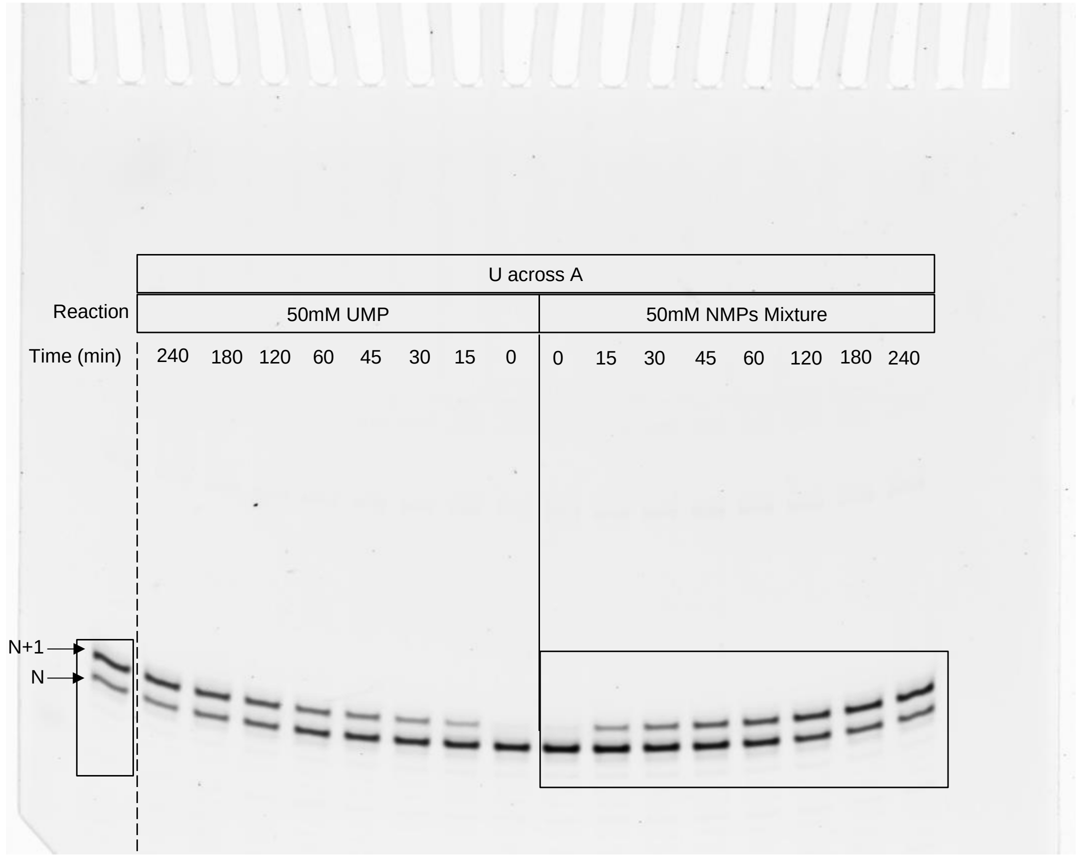


**Figure S18**: Box represents the cropped part of $U_A$ reaction shown in Figure 1(iv)A in the main text.

#### HPLC Hydrolysis Profiles of ImpNs

ImpNs were dissolved in ultrapure water (pH-7±0.05). Their concentrations were determined and equal amounts were loaded on to the HPLC (Infinity Series 1260HPLC, Agilent Technologies). The activated and non-activated molecules were separated on DNAPac PA 200 column (Thermo Scientific). A previously standardized method was used to separate activated nucleotide from the non-activated spent monomer[1].

Table S1: Hydrolysis profiles of ImpNs at different time points

| Stock | % Species at t=0 | | % Species at t=x | | |
|---|---|---|---|---|---|
| | ImpN | NMP | x (min)# | ImpN | NMP |
| **ImpA** | 96.6 | 3.4 | 30 | 88.0 | 12.0 |
| **ImpG** | 77.7 | 22.3* | 30 | 68.4 | 31.6 |
| **ImpC** | 86.7 | 13.3* | 5 | 85.3 | 14.7 |
| **ImpU** | 72.1 | 27.9* | 60 | 72.0 | 28.0 |

* Although there is hydrolysis observed in ImpG, ImpC and ImpU, it is pertinent to note that only 2.2mM, 1.3mM and 2.7mM of hydrolysed GMP, CMP, and UMP, respectively, made itto the actual reaction from these ImpN stocks. This concentration is much lesser when compared to the concentration of the co-solutes that were added externally to the reaction mixtures.

### This t reflects the reaction timeline used for the calculation of the initial reactions rates for the various reactions studied.

**References**:


1. Dagar, S., Sarkar, S. & Rajamani, S. Geochemical influences on nonenzymatic oligomerization of prebiotically relevant cyclic nucleotides. *RNA* **26**, 756–769 (2020).
